# The theoretical molecular weight of NaYF_4_:RE upconversion nanoparticles

**DOI:** 10.1101/114744

**Authors:** Lewis E. Mackenzie, Jack A. Goode, Alexandre Vakurov, Padmaja P. Nampi, Sikha Saha, Gin Jose, Paul A. Millner

**Author notes:** **Corresponding author:** Lewis MacKenzie.

## Abstract

Upconversion nanoparticles (UCNPs) are utilized extensively for biomedical imaging, sensing, and therapeutic applications, yet the molecular weight of UCNPs has not previously been reported. We present a theory based upon the crystal structure of UCNPs to estimate the molecular weight of UCNPs: enabling insight into UCNP molecular weight for the first time. We estimate the theoretical molecular weight of various UCNPs reported in the literature, predicting that spherical NaYF4 UCNPs ~ 10 nm in diameter will be ~1 MDa (i.e. 10^6^ g/mol), whereas UCNPs ~ 45 nm in diameter will be ~100 MDa (i.e. 10^8^ g/mol). We also predict that hexagonal crystal phase UCNPs will be of greater molecular weight than cubic crystal phase UCNPs. Additionally we find that a Gaussian UCNP diameter distribution will correspond to a lognormal UCNP molecular weight distribution. Our approach could potentially be generalised to predict the molecular weight of other arbitrary crystalline nanoparticles: as such, we provide standalone graphic user interfaces to calculate the molecular weight both UCNPs and arbitrary crystalline nanoparticles. We expect knowledge of UCNP molecular weight to be of wide utility in biomedical applications where reporting UCNP quantity in absolute numbers or molarity will be beneficial for inter-study comparison and repeatability.

## Introduction

Photonic upconversion nanoparticles (UCNPs) have garnered widespread scientific interest due to their unique near infra-red (NIR) excitation and visible luminescence properties; a process known as photonic upconversion. UCNPs are inorganic crystalline nanostructures (typically NaYF4) co-doped with rare-earth (RE) ions, (e.g. Yb^3+^, Er^3+^, Gd^3+^); hereby referred to in general terms as NaYF_4_:RE UCNPs. The RE ions act as sensitizers and emitters for photonic upconversion of multiple infra-red photons, resulting in visible luminescence emission. UCNP emission is highly stable,^1^ with no photo bleaching, and a relatively long luminescence emission lifetime ranging from hundreds of microseconds to a few milliseconds.^2,3^ NIR excitation via upconversion is highly advantageous for biomedical applications, where ultraviolet or visible excitation of fluorophores (e.g. dyes, proteins, or quantum dots) is normally required, with the associated challenges of photo-bleaching and photo-toxicity. Interactions between nearby molecules and the UCNPs crystal structure enables molecular biosensing via luminescence resonance energy transfer (LRET) between UCNPs and molecules in proximity to them. ^4–10^ As such, UCNPs have found wide utility in biomedical applications, including as imaging contrast labels in cellulo, in vivo, and ex vivo^5,11–19^; as biosensors for detection of antibiotics^20^ and toxins in food^21–23^; as biosensors to measure biomarkers in biological fluids (e.g. whole blood, serum, urine),^6–8,24–26^ and as therapeutic agents, against targets such as cancer cells.^27,28^ Additionally UCNPs have been applied to nanoscale thermometry^29,30^ and photovoltaic applications.^31,32^ However, to date, the molecular weight of UCNPs has not been reported: as such, both the molarity of UCNPs in solution, and the absolute number of UCNPs in a sample has been unknown.

The lack of molecular weight information for UCNPs is a considerable shortcoming in biomedical applications of UCNPs, where precise quantification of UCNP concentration would be highly beneficial for informing of dosage of UCNPs studies, as well as aiding inter-study comparison. Additionally, quantification of UCNP molarity and absolute number of UCNPs would be highly beneficial when constructing biosensors where the ratio of UCNPs compared to other molecules, e.g. antibodies^6–8,25^ or oligonucleotides,^33^ is important for informing biosensor design.

The lack of information on UCNP molecular weight is likely due to lack of experimental techniques capable of measuring the molecular weight of large macromolecules such as UCNPs. Using the theory we present in this paper, we predict that the molecular weight of NaYF_4_:RE UCNPs will range from a few mega Daltons (MDa) (i.e. 10^6^ g/mol) for exceptionally small UCNPs (~10 nm in diameter), to > 100 MDa (for UCNPs with a more typical diameter of ~45 nm). This large molecular weight range is well beyond the measurement limits both mass spectrometry and sedimentation velocity analytical ultracentrifugation (svUAC), which are limited to < 40 kDa and < 5 MDa respectively.^34^ Despite this limitation, we attempted to employ svAUC to estimate the molecular weight of UCNPs ~30 nm in diameter (corresponding to a molecular weight of ~40 MDa), but reliable results were not obtained (see the Discussion section and supplementary material).

In this study, we present a theoretical method, based upon the extensively studied and empirically proven theory of crystallography and UCNP structure, to calculate the molecular weight UCNPs, accounting for UCNP composition and morphology. In brief, the crystalline structure of UCNPs is quantified by transmission electron microscopy (TEM), and x-ray diffraction (XRD) experiments. From this information, the total atomic weight within a single NaYF_4_:RE unit cell, and the total number of unit cells within a UCNP can be calculated. Thus, the theoretical molecular weight of UCNPs can be calculated by summing up the total molecular weight contained within all unit cells in a UCNP.

We anticipate that this theoretical framework could be extended to crystalline nanoparticles of arbitrary morphology and composition, provided that the crystalline structure of such nanoparticles are known. As such, we also provide two stand-alone graphical user interfaces (GUIs) for simple calculation of the molecular weight of both NaYF_4_:RE UCNPs and arbitrary crystalline nanoparticles. Knowledge of UCNP molecular weight will likely be highly beneficial for quantification of UCNP concentration in biomedical applications.

## Theory

### Crystalline structure and photonic upconversion properties of UCNPs

The key to understanding both the optical properties of UCNPs and their molecular weight lies in the crystalline structure of UCNPs. UCNPs are a crystal lattice made up of repeating crystal unit cells of NaYF4, with a fraction of Y^3+^ ions selectively replaced by RE dopants (see Figure 1). In UCNPs, photonic upconversion is enabled by the absorption of two or more near-infrared photons, which, via excitation of several long-lived metastable electron states, and subsequent non-radiative multi-phonon and radiative relaxation, produces luminescence emission at visible wavelengths (see Figure 2). Efficient upconversion requires a crystalline host lattice, which is doped with multiple different lanthanide ions (typically Yb^3+^ and Er^3+^), where one lanthanide ion acts as a photosensitizer (typically Yb^3+^) and acts as a photonic emitter (typically Er^3+^).^35^ Although many different combinations of lattice and RE dopants have been explored,^36^ the combination of Yb^3+^ and Er^3+^ ions in a NaYF4 host lattice has been found to provide high upconversion efficiency, and as such is commonly used for UCNPs.^37,38^ Figure 2 shows an exemplar upconversion emission spectrum of NaYF4:Yb,Er cubic UCNPs (20% Yb^3+^, 2% Er^3+^) and the corresponding Jablonski diagram for 39 upconversion.

**Figure 1.**
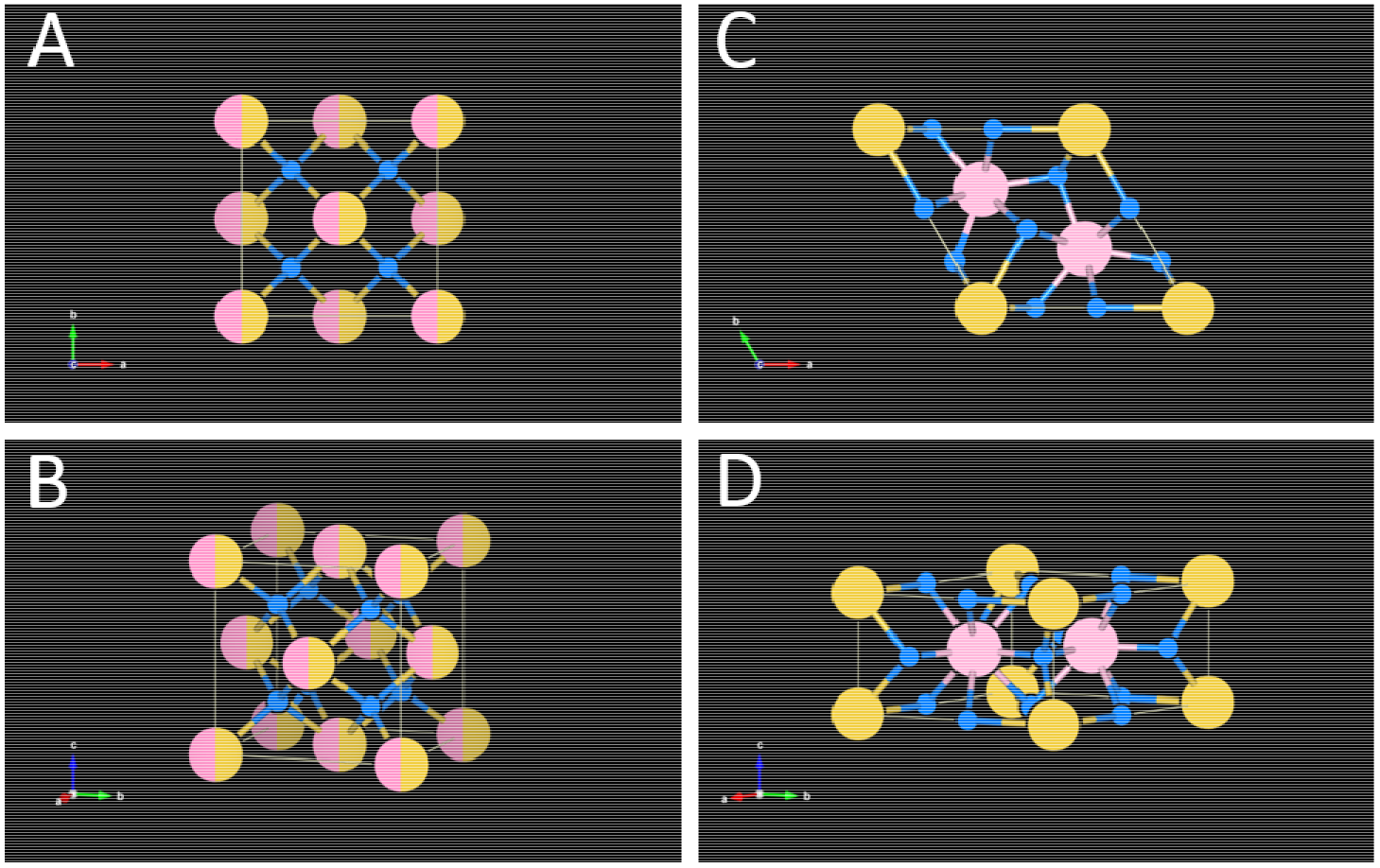
NaYF_4_:RE UCNPs unit cell structures. Colour key: Na^+^ ions are yellow; Y^3+^ and RE^3+^ dopant ions are pink; F^-^ ions are smaller and blue. **(A, B)** Cubic lattice unit cell structure. Sites that are randomly occupied by both Na^+^ and RE^3+^ are depicted as both pink and yellow. **(C, D)** Hexagonal lattice unit cell structure. This Figure is based upon data from Kramer et al., (2004)^42^ and Wang et al., (2010).^40^ Diagrams created with the open-source software package VESTA.^43^

**Figure 2.**
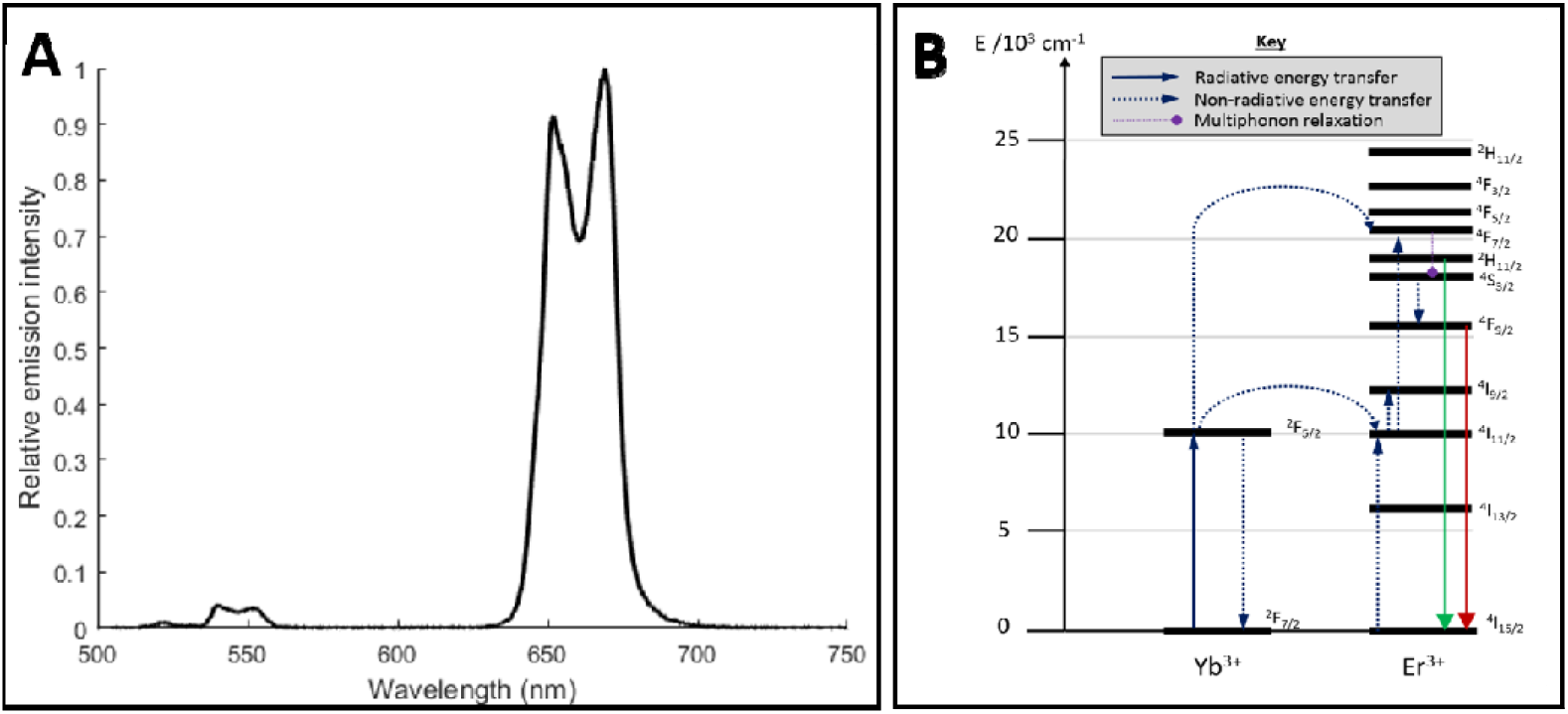
The upconversion emission of UCNPs. **(A)** Emission of a 1 mg/mL of NaYF4:Yb,Er UCNPs (20% Yb and 2% Er) suspended in ultra-pure water (see supplementary material for UCNP synthesis details). **(B)** Corresponding Jablonski diagram depicting the upconversion process (based upon Heer et al., (2004)).^39^

NaYF_4_:RE unit cells are either a cubic or a hexagonal crystal lattice arrangement (see Figure 1). In the face-centred cubic lattice arrangement (Na_2_Y_2_F_8_), high-symmetry cation sites are formed, and are randomly occupied by either Na^+^ or RE^3+^ ions (see Figure 1a), and Y^3+^ ions are substituted for other RE^3+^ ions, enabling photonic upconversion. In hexagonal unit cells (Na_1.5_Y_1.5_F_6_), there are two relatively low-symmetry cation sites, which contain either Na^+^ or RE^3+^ ions (see Figure 1b).^40^ Characterisation of UCNP unit cells is typically conducted by XRD measurements. Several studies have reported the crystal lattice parameters associated with cubic and hexagonal NaYF_4_:RE UCNPs: these are summarised in Table 1. Wang et al., (2010)^40^ report unit cell parameters for cubic (α phase) and hexagonal (β phase) unit NaYF_4_:RE unit cell configurations (see Figure 1). The arrangement of ions within unit cells influences the crystal lattice parameters, consequently changing photonic properties, such as upconversion quantum efficiency.^40^

Synthesis of NaYF_4_:RE UCNPs typically creates pseudo-spherical UCNPs with a range of diameters. For example, Sikora et al., (2013) report a Gaussian diameter distribution of UCNPs, ranging between 15 – 70 nm.^14^ and Haro-González et al. (2013)^41^ report a Gaussian diameter distribution of UCNPs UCNPs ranging from ~10 – 50 nm in diameter (see Table 1).

**Table 1.** Crystal lattice parameters of NaYF_4_:RE UCNPs reported in the literature.

| Study | NaYF<sub>4</sub> RE dopant composition (%) | UCNP lattice structure | a (Å) | c (Å) | Mean UCNP diameter (nm) | UCNP diameter range (nm) |
| --- | --- | --- | --- | --- | --- | --- |
| Sikora et al., (2013).<sup>14</sup> | 30% Yb<sup>3+</sup>, 2% Er<sup>3+</sup> | Cubic | 5.51 | - | ~ 30 | 15 -70 |
| Cao et al., 2010.<sup>15</sup> | 20% Yb<sup>3+</sup>, 2% Er<sup>3+</sup> | Hexagonal | 5.960 | 3.510 | 33 ± 1 nm | 32 - 34 nm |
| Wang et al., 2010.<sup>40</sup> | 18% Yb<sup>3+</sup>, 2% Er<sup>3+</sup> | Hexagonal | 5.96 | 3.53 | Not reported | Not reported |
| Wang et al., 2010.<sup>40</sup> | 18% Yb<sup>3+</sup>, 2% Er<sup>3+</sup>, 60% Gd<sup>3+</sup> | Hexagonal | 6.02 | 3.60 | Not reported | Not reported |

### Estimating the number of unit cells in a UCNP

For the purposes of this study, we assume UCNPs to be spherical, with volume (*V_UCNP_*) described by:

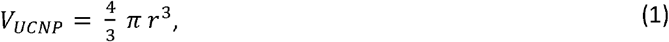

where *r* is the radius of the UCNP. Note that non-spherical UCNP morphologies can be incorporated by modifying Equation 1 appropriately. If the UCNP consists of cubic unit cells, then the volume of an individual cubic unit cell (*uV_cubic_*) is given by:

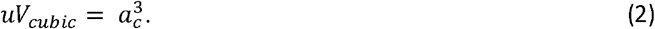

If the UCNP consists of hexagonal unit cells, volume of a hexagonal unit cell (*uV_hexagonal_*) is given by:

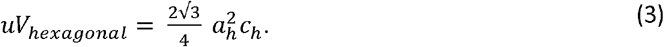

Where *a_h_* and *c_h_* are lattice parameters describing hexagonal unit cells. Thus, the number of unit cells in a UCNP (i.e. *uN_cubic_* or *uN_hexagonal_*) can be estimated by:

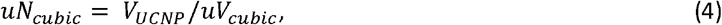

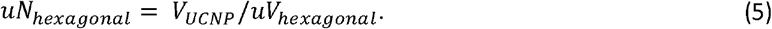

This calculation assumes the effects of crystal dislocations and rounding error in the total number of unit cells to be negligible. Further, we assume that UCNPs are composed of 100% cubic or hexagonal unit cells because, to the best of our knowledge, hybrid crystal phase UCNPs have not been reported.

### Estimating the total atomic weight within a single unit cell

Assuming no RE dopants, the atomic weight of a single cubic NaYF4 *uAW_cubic_* or hexagonal NaYF4 unit cell *uW_hex_* is described by:

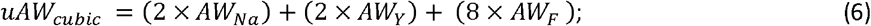

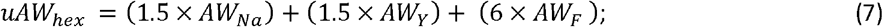

where *AW_Na_*, *AW_Y_*, and *AW_F_* are the atomic weight (Da or g/mol) of Sodium, Yttrium, and Fluorine respectively (see Table S1). We assume any mass difference due loss of electrons due to ionisation to be negligible. If RE dopant ions are added during UCNP synthesis, then a fraction of Y^3+^ ions are substituted for RE^3+^ dopant ions, altering the average atomic weight of unit cells within UCNPs. This RE doping can be accounted for by defining a total additive factor (*af*):

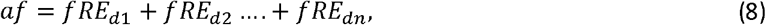

where *RE_d1_*, *fRE_d2_*, … *fRE_dn_* is the fractional percentage of an arbitrary number of RE dopants. The total additive factor is a numeric value ranging between 0 and 1, representing the theoretical extremes of 0% and 100% Y substitution respectively. Thus, total the atomic mass contained withi a single cubic or hexagonal unit cell with RE dopants is be calculated by:

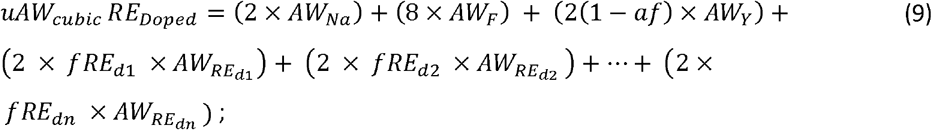

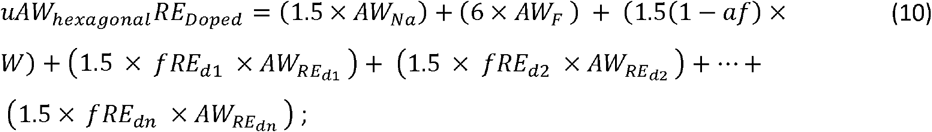

where *uAW_cubic_ RE_Doped_* and *uAW_hexagonal_ RE_Doped_* are the average atomic weight of RE doped cubic and hexagonal unit cells, respectively.

### Estimating the theoretical molecular weight of a UCNP

Once the total number of unit cells within a UCNP (*uN*) and the total atomic weight (*uAW*) within each individual unit cell are estimated, the theoretical molecular weight of a cubic lattice UCNP *MW_cubic_* can be estimated by summing the atomic weight contributions from all unit cells:

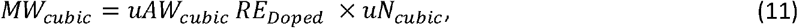

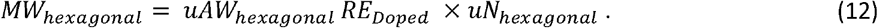

From Equations 4, 5, 11, and 12, it can be seen that the molecular weight of UCNPs scales proportionally to volume, thus spherical UCNPs molecular weight will scale proportionally to the cube of UCNP radius.

## Methods

### Molecular weight predictions for cubic and hexagonal NaYF_4_:RE UCNPs

Using the theory presented in Sections 2.4 – 2.6, the theoretical molecular weight of hexagonal and cubic lattice NaYF4 UCNPs were calculated, assuming the following typical unit cell lattice parameters: cubic: *a* = 5.51 Å; hexagonal: *a* = 5.91 Å, *c* = 3.53 Å; (see Table 1).

### The effect of RE doping on theoretical molecular weight

The effect of RE doping was investigated by using the theory presented in Sections 2.4 – 2.6 to calculate the theoretical molecular weight of NaYF_4_:RE UCNPs incorporating various concentrations of Yb^3+^ and Er^3+^ dopant ions. We assume that UCNP lattice parameters will remain constant, neglecting the unit cell contraction effect demonstrated by Wang et al. (2010),^40^ where UCNP unit cell lattice parameters are altered when the concentration of RE dopants is increased.^40^

### The theoretical molecular weight of UCNPs reported in the literature

The theoretical molecular weight of various NaYF_4_:RE UCNPs reported in the literature was calculated by incorporating various lattice parameters and mol% of RE dopants from the literature into the theory presented in Sections 2.4 – 2.6.

### UCNP diameter distribution vs. theoretical molecular weight distribution

UCNP synthesis typically produces a Gaussian distribution of UCNPs diameters. To investigate how such a distribution of UCNP diameters affects the distribution of theoretical UCNP molecular weights, the Gaussian diameter distribution data for a single batch of NaYF4:Yb,Er UCNPs was reproduced from data presented in Sikora et al (2013).^14^ The theoretical molecular weight for each UCNP diameter in this distribution was calculated by the theory presented in Sections 2.4 – 2.6. Gaussian fits to the data were calculated by using non-linear least squares fitting in MATLAB (MATLAB 2016a, MathWorks).

### Stand-alone GUIs for calculation of nanoparticle theoretical molecular weight

Two stand-alone executable graphic user interfaces (GUIs) were created in MATLAB to enable rapid calculation of UCNP theoretical molecular weight. Each GUI incorporates different features and assumptions. The first, more simple, GUI was developed to enable other researchers to calculate the theoretical molecular weight of spherical NaYF_4_:RE UCNPs for a user-defined nanoparticle size range. The second, more powerful, GUI was designed to enable users to estimate the theoretical molecular weight of crystalline nanoparticles with arbitrary nanoparticle geometry; arbitrary lattice parameters; and arbitrary elemental composition, across a user-defined range of characteristic nanoparticle sizes. Additional technical information for both GUIs is provided in the supplementary material section. The stand-alone GUIs developed are shown in supplemental Figures S1 and S2. These GUIs are freely available from the University of Leeds Research Data Depository and are attributed with their own citable DOI (https://doi.org/10.5518/173.^44^

## Results

### Theoretical molecular weight of cubic and hexagonal NaYF_4_:RE UCNPs

Hexagonal lattice UCNPs have a greater theoretical molecular weight than cubic lattice UCNPs (see Figure 3); this is due to the lower volume of hexagonal unit cells, and correspondingly higher density of hexagonal lattice UCNPs. Additionally, because molecular weight scales to UCNP volume, relatively small changes in UCNP diameter increased molecular weight considerably: e.g. a 20 nm cubic UCNP has a molecular weight of ~10 MDa, whereas a 30 nm UCNP has a molecular weight of > 30 MDa (an increase of 20 MDa for a 5 nm change in UCNP diameter).

**Figure 3.**
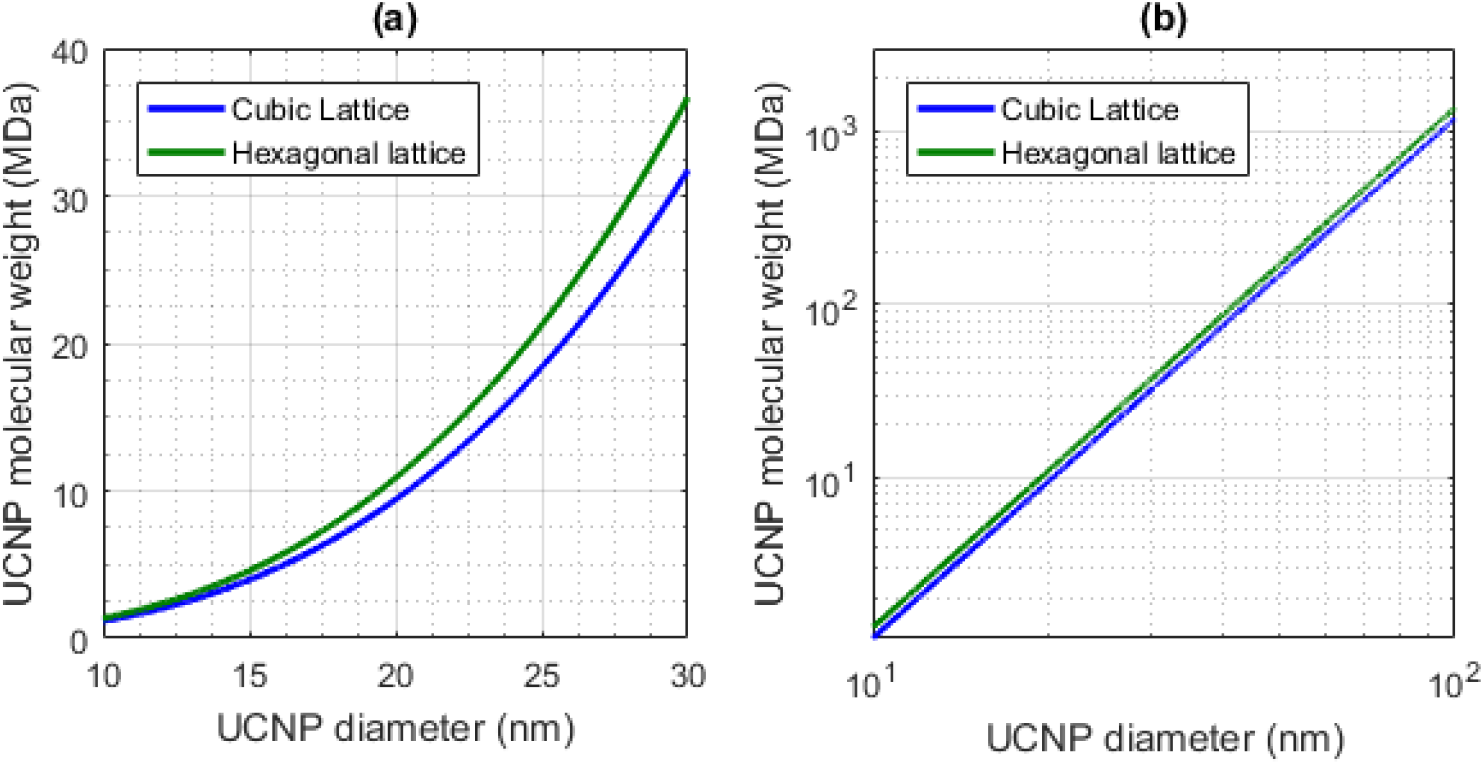
Diameter versus theoretical molecular weight for hexagonal and cubic NaYF4 UCNPs (green and blue respectively). **(a)** UCNP diameter vs. molecular weight on a standard x-axis. **(b)** The same data plotted with a logarithmic scale. Lattice parameters were assumed to be: a = 5.51 Å for cubic UCNPs; a, c = 5.91 Å and 3.53 Å for hexagonal UCNPs.

### The effect of RE doping on UCNP molecular weight

Increasing Yb^3+^ or Er^3+^ dopant % increased the theoretical molecular weight of UCNPs (see Figure 4) because Yb^3+^ and Er^3+^ have a greater atomic mass than Y^3+^. However, the difference in theoretical molecular weight between UCNPs doped with Yb^3+^ and Er^3+^ was relatively small to the small difference between the atomic weight of Yb^3+^ and Er^3+^ (173.054 and 167.259 g/mol respectively, see Table S1). Hexagonal lattice UCNPs show a slightly higher increase in theoretical molecular weight for a given dopant concentration than cubic lattice UCNPs because hexagonal lattice UCNPs have a greater unit cell density compared to their cubic counterparts.

**Figure 4.**
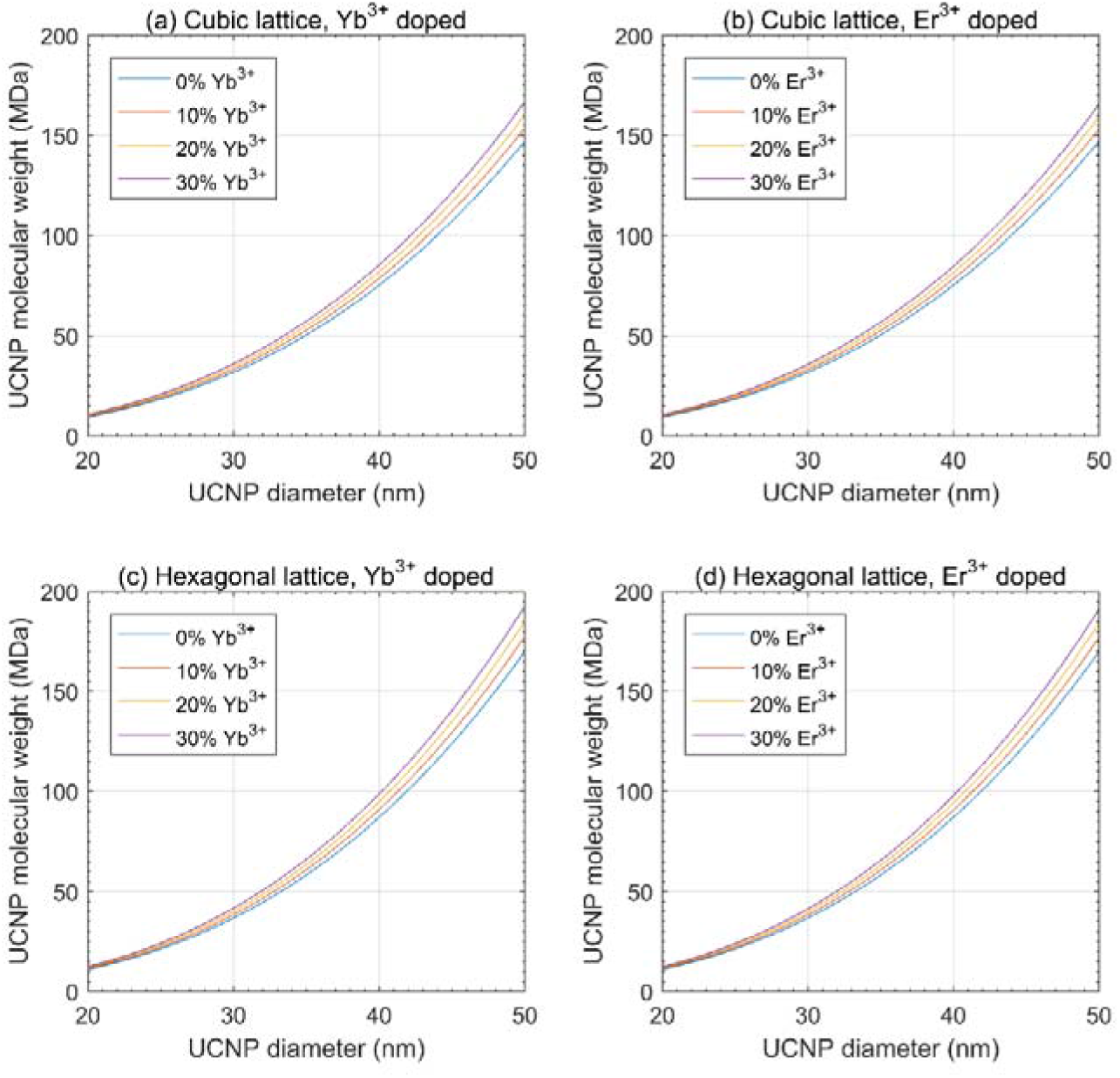
The effect of RE doping on theoretical UCNP molecular weight. **(a, b)** theoretical molecular weight vs. RE dopant mol% for cubic lattice UCNPs. **(c, d)** theoretical molecular weight vs. RE dopant mol% for cubic lattice UCNPs. Calculations assume that lattice parameters are a = 5.51 Å for cubic lattice UCNPs, a = 5.91 Å; c = 3.53 Å for hexagonal lattice UCNPs, and that lattice parameters are independent of dopant mol%.

### The theoretical molecular weight of NaYF_4_:RE UCNPs reported in the literature

The theoretical molecular weight of various NaYF_4_:RE UCNPs reported in the literature are shown in Figure 5.

**Figure 5.**
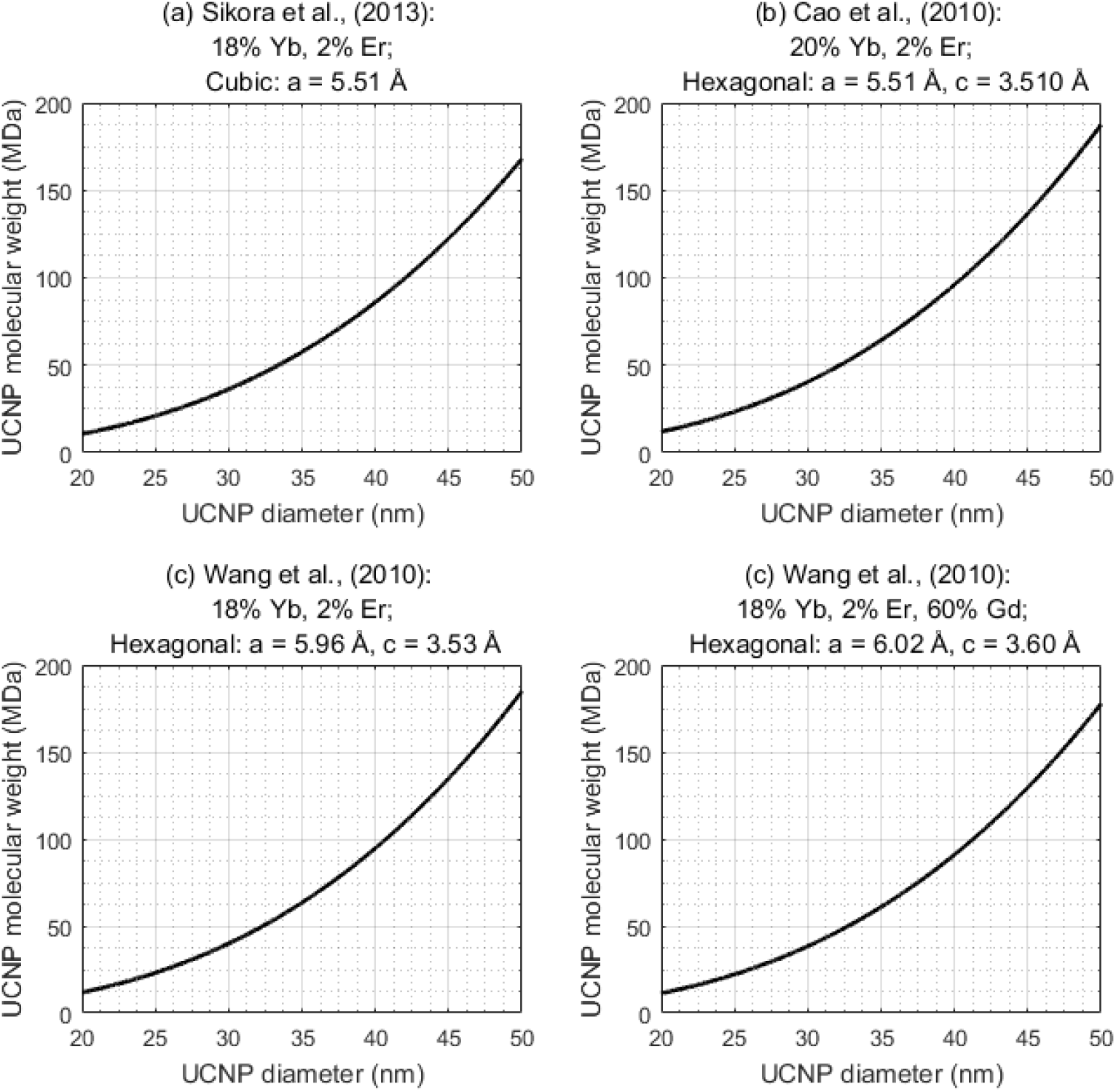
Theoretical molecular weight of various UCNPs reported in the literature. **(a)** Sikora et al., (2013).^14^ **(b)** Cao et al., (2010).^15^ **(c, d)** Wang et al., (2010).^40^

### UCNP diameter distribution vs. theoretical molecular weight distribution

The UCNP diameter distribution data from Sikora et al., (2013)^14^ was well-fitted by a Gaussian distribution (R^2^ = 0.96) (see Figure 6a). The corresponding theoretical molecular weight distribution (shown in Figure 6b), demonstrates the exponential relation between UCNP diameter and UCNP molecular weight distribution. Plotted on a logarithmic x-axis scale (Figure 6c), the resulting molecular weight distribution was well fitted by a Gaussian distribution (R^2^ = 0.98), indicating that the molecular weight distribution corresponding to a Gaussian diameter distribution is lognormal.

**Figure 6.**
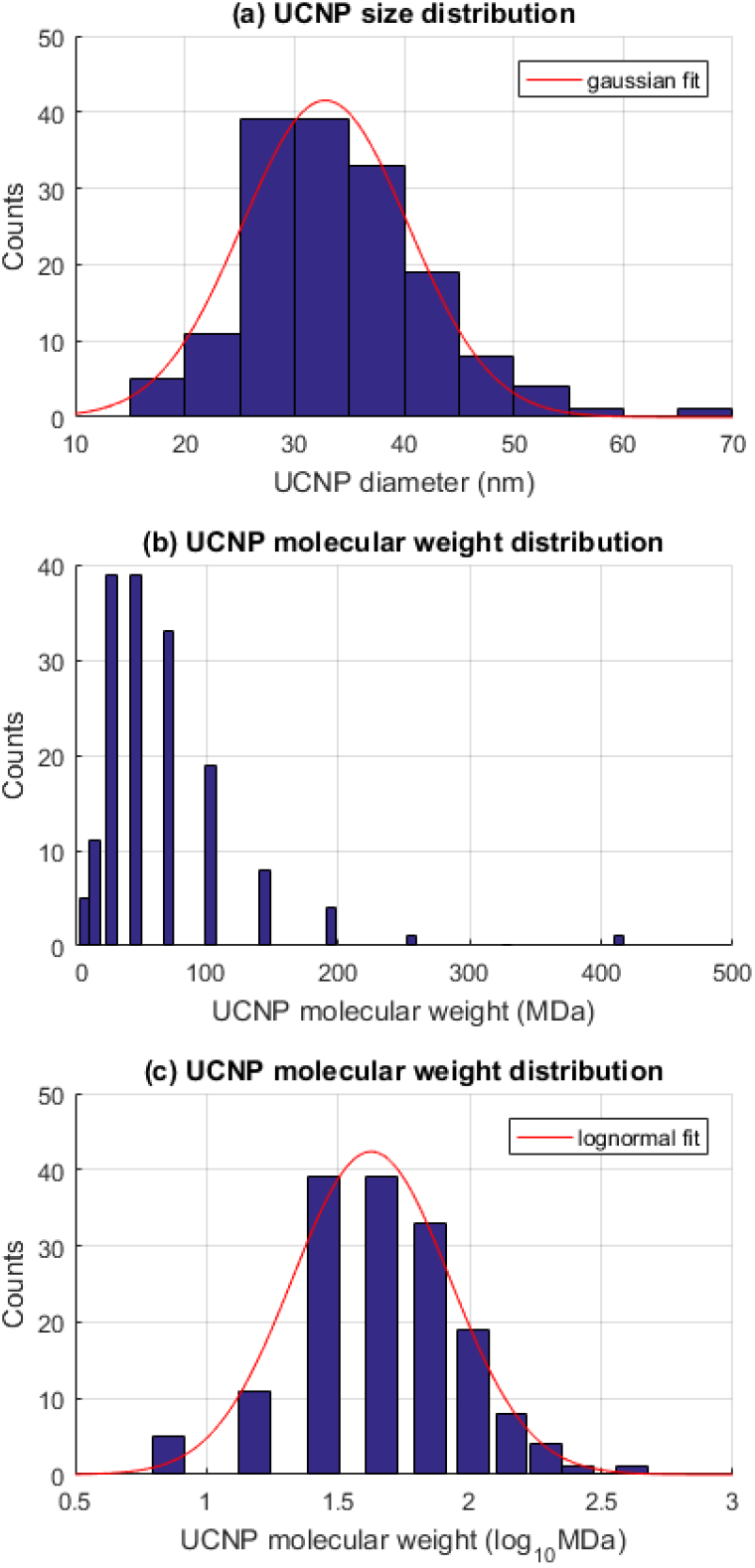
Gaussian UCNP diameter distributions give arise to lognormal distribution of theoretical molecular weights. **(a)** A Gaussian diameter distribution of UCNPs is well described by a normal distribution (R^2^ = 0.96). **(b)** The corresponding theoretical molecular weight distribution of UCNPs on a linear molecular weight scale. **(c)** The molecular weight distribution on a logarithmic x-axis is well fitted by a lognormal distribution (R^2^ = 0.98).

## Discussion

We have provided a theory to estimate the molecular weight of UCNPs. Our theory is required because, to the best of our knowledge, there are no experimental techniques measuring the molecular weight of UCNPs, which we predict will be > 5 MDa for UCNPs ~15 nm in diameter, and 100 MDa for UCNPs ~45 nm in diameter. Mass spectrometry is limited to molecules < 40 kDa, and svAUC is limited to measurements of macromolecules < 5 MDa.^34^ Our theory predicts that UCNPs with a molecular weight of < 5 MDa would be < 15 nm in diameter. To the best of our knowledge monodisperse synthesis of such small UCNPs has not been reported in the literature.

Despite the aforementioned challenges of experimental verification, we attempted svAUC measurements of UCNPs, because successful svAUC studies of other types of nanoparticles (e.g. SiO2 nanoparticles) with unknown molecular weight have been reported.^45,46^ If accurate svAUC measurements of UCNPs could be made, then UCNP molecular weight could potentially be calculated by the theory described by Carney et al., (2011),^34^ which is based upon accurate quantification of sedimentation and diffusion coefficients from svAUC measurements, and which has been verified for gold nanoparticles ~2 MDa in molecular weight. The full details of the method of our svAUC experiment are provided in the supplementary information. However, our svAUC experiment studying UCNPs was not successful. In brief, our avAUC results showed that the UCNPs (diameter = 32 ± 5 nm, average theoretical molecular weight of ~ 43 MDa) sedimented very rapidly, even at low centrifuge rotor speeds (3,000 rpm), limiting the amount of useable data. At higher rotor speeds UCNPs sedimented too rapidly for data collection. When the recovered sedimentation coefficient was extrapolated to zero sample concentration, a negative sedimentation coefficient was returned. Additionally, UCNPs were observed to diffuse considerably, further complicating AUC experiments. This unusual behaviour is not typical of the nanoclusters and gold nanoparticles used to demonstrated the molecular weight estimation technique described by Carney et al., (2011),^34^ and as such UCNP molecular weight could not be estimated by svAUC. The challenges associated with svAUC measurement of UCNPs serve to further highlight the need for a method to estimate the molecular weight of UCNPs theoretically.

Although it has not been possible for us to experimentally validate our estimates of UCNP molecular weight, it may be possible in future to verify some limited predictions of our theory. For example, it may be possible to measure the difference in bulk densities of cubic and hexagonal UCNPs and compare this with predictions from our theory. However, we could not attempt this measurement because we did not have access to the high temperature crucible equipment required for hexagonal UCNP synthesis.^40^

Despite this lack of current and direct experimental verification, we can be reasonably confident in the accuracy of our theory because it stems directly from the theory of crystallography, which has been a subject of intense study in the past century,^47^ combined with empirical measurements of UCNP crystal structure.

Our method to calculate the theoretical molecular weight of NaYF_4_:RE UCNPs relies on two basic assumptions: 1. that UCNPs are crystals of homogenous elemental composition and unit cell phase, and 2. that the lattice parameters and diameter data utilized is accurate. These assumptions can be verified by TEM and XRD measurements of UCNP crystal structure. Ensuring accurate lattice parameters is particularly important when estimating the molecular weight of UCNPs with arbitrarily large dopant concentrations. For example, Wang et al., (2010)^40^ experimentally demonstrated that by doping a hexagonal phase NaYF4:Yb,Er UCNP (18% Yb, 2% Er) with increasing concentrations of Gd^3+^ increases the lattice parameters of the UCNP significantly, resulting in an increased unit cell volume. Thus, because of this dependence of lattice parameter on RE dopant percentage, our estimations of UCNP molecular weight in Figure 3 may be an over-estimation on true values if lattice parameters are not independently verified for each RE dopant concentration of interest. UCNP volume/morphology also influences theoretical UCNP molecular weight. We recommend using TEM to directly quantify UCNP morphology with limited assumptions. Other techniques such as such as dynamic light scattering (DLS) and nanoparticle tracking analysis can be used to estimate the equivalent hydrodynamic radius of nanoparticles but incorporate various assumptions into calculations.^46,48^ As such, direct TEM imaging of UCNPs is preferable to ensure theoretical molecular weight is as accurate as possible. In this study we assumed UCNPs are perfectly spherical, but our method could be trivially adapted for arbitrary nanoparticle geometries; e.g. rods,^40,49^ triangular,^50^ or prism-shaped^51^ nanoparticles, and for nanoparticles of varying crystalline composition. The extension of our technique to arbitrary geometries, arbitrary crystal lattice parameters, and arbitrary elemental composition is demonstrated by the development and application of an advanced GUI incorporating all of these variables (see Figure S2). Our theory does not account for any dislocations in the regular UCNP crystal structure. Instead we assume the influence of any such dislocations to be negligible compared to the molecular weight of whole UCNPs. Our theory also does not account for any surface functionalisation with amorphous layers or other molecules. Thus the molecular weight of UCNPs modified by addition of a silica^8,35,52^ or calcium fluoride^53^ shell coating will be greater than the theoretical molecular weight estimated by our technique.

It should be noted that a simple theory for estimation of the molecular weight of a single homogenous gold nanoparticle based upon bulk density of materials was proposed by Lewis et al. (2006).^54^ However, this simple theory did not account for crystalline unit cell parameters or elemental doping. Further, their theory was not extended to describe the molecular weight distributions of a population of nanoparticles. Our results demonstrate that a Gaussian distribution of UCNP diameters corresponds to a lognormal distribution in molecular weight (as shown in Figure 6). Mathematically, it is reasonable to expect similar logarithmic relations between UCNP diameter and molecular weight for arbitrary diameter distributions. Such molecular weight distributions may of consequence when studying behaviour of UCNP populations, because minor outliers in UCNP diameter will be extreme outliers in terms of molecular weight.

Estimation of molecular weight of NaYF_4_:RE UCNPs will likely be of utility in various applications, particularly in biomedical imaging, biosensing, and therapeutics. Knowledge of UCNP molecular weight will likely be of great utility in studies where UCNP surfaces are functionalised with additional molecules, e.g. antibodies ^6–8,25^ or oligonucleotides,^33^ because If the molecular weight of UCNPs is known, then the molar concentrations of substances in the functionalisation processes can be determined. When combined with estimation of UCNP surface area, this could inform the UCNP functionalisation for biosensing applications. Knowledge of UCNP molecular weight would also be beneficial in the processing of particles for downstream applications. In particular, steps taken to functionalise the nanoparticles may require separation procedures to remove unreacted moieties or unwanted reactants. If the molecular weight of UCNPs were known, then it may be beneficial for the optimisation of conjugation stoichiometry, which can be concentration dependant; the reaction rates of UCNPs will be heavily influenced by their molecular weight; thus a greater understanding of their molecular weight may increase the knowledge of thermodynamic properties of UCNP systems. This is particularly important when considering the use of bioreceptors with UCNPs where the mass of the particle may affect the binding kinetics of the UCNP-receptor construct.

The molecular weight of UCNPs will also be of interest in the study of cytotoxicity, biodistribution, cellular uptake, metabolism, and excretion of UCNPs in biological systems.^12,14^ Currently, it is extremely challenging to compare the results from various imaging and therapeutic studies because UCNP concentration is reported as weight of UCNPs per volume of aqueous media (i.e. mg/mL or similar).^12^ This is a crude measure which does not quantify number of UCNPs in a given sample. For example, nanoparticles can induce membrane damage^55^ and initiate apoptosis (programmed cell-death).^56,57^ Reporting the molar concentration of UCNPs would help assessment of UCNP cytotoxic effects. A standardised protocol based on molecular weight of UCNPs would help assessment of accumulation of UCNPs in vivo and their clearance time from organs^13^ or tumours.^53^ Reporting the molar concentration of UCNP composites may also help to develop highly-localised targeted delivery of therapeutic drugs to the required sites in the body, leading to better controlled targeted photodynamic therapy,^27^ and potential improvements in targeted drug delivery.^16^

## Conclusions

We have provided a method to estimate the theoretical molecular weight of UCNPs. This theory is based upon UCNP crystal parameters which can be measured for batches of UCNPs by TEM and XRD techniques. The theory presented here is generalizable to other crystalline nanoparticles where the relevant crystalline lattice parameters are known, i.e. nanoparticle unit cell elemental composition, unit cell size parameters, and nanoparticle morphology. To enhance dissemination of our theory we provide two stand-alone GUIs for calculation of the molecular weight of both UCNPs and arbitrary crystalline nanoparticles respectively. We could not, however, experimentally verify our predictions of UCNP molecular weight with mass spectrometry or svAUC. We did attempt svAUC experiments but could not recover reliable svAUC data because UCNPs were observed to sediment and diffuse rapidly. Nevertheless, our theory provides some key predications about the molecular weight of UCNPs. Firstly, that the theoretical molecular weight of UCNPs scales with volume of the nanoparticle. As an example, we predict that a spherical UCNP ~10 nm diameter will have a molecular weight of ~1 MDa (10^6^ g/mol), whereas a UCNP ~ 45 nm in diameter will be ~100 MDa (10^8^ g/mol). From this relation, we find that a Gaussian distribution of nanoparticle diameters corresponds to a lognormal distribution of UCNPs molecular weights, and that a small change in UCNP diameter distribution can potentially represent a large change in overall UCNP molecular weight. We also report that Hexagonal crystal lattice phase UCNPs will be of greater molecular weight than cubic lattice phase UCNPs, and that increasing RE dopant % will increase UCNP molecular weight.

We expect that the knowledge of UCNP molecular weight will be of utility in a wide variety of biomedical applications, as UCNP concentrations can now be reported in terms of molarity or absolute number of UCNPs instead of the relatively crude measure of sample weight. This will likely aid inter-study comparison of both UCNP dosage and improve methods for creating UCNP biosensors.

## Acknowledgements

The authors would like to extend a special thanks to Amy Barker (School of Molecular and Cellular Biology, University of Leeds) for her technical assistance and expertise in conducting and analysing AUC experiments. We are also grateful to Professor Peter Stockley (School of Molecular and Cellular Biology, University of Leeds) for granting access to the AUC facilities.

## Funding Acknowledgements

- L.E. MacKenzie was supported by a grant from the Biotechnology and Biological Sciences Research Council Tools and Development Resources Fund (BBSRC TDRF) (BB/N021398/1).
- J.A. Good is supported by a grant from the Medical Research Council (MRC) (MR/N029976/1).
- A. Vakurov was supported by a grant from the Natural Environment Research Council (NERC).(NE/N007581/1).
- Padmaja P. Nampi is supported by a European Commission Marie Skłodowska-Curie Individual Fellowship for Experienced Researchers (H2020-MSCA-IF-2015).

## Competing financial interest statement

The authors declare no competing financial interests

## Author contributions statement

- L.E.M and J.A.G. conceived the research concept and wrote the manuscript.
- L.E.M. performed all calculations, provided Figures 2, 3, 4, 5, 6, and created the stand-alone GUIs.
- J.A.G. provided Figure 1.
- A.V, P.P.N, S.S, G.J., and P.M, contributed to and reviewed the manuscript.

